# Maternal age, obesity and hyperglycaemia are associated with a delay in preimplantation development in a mouse model of type 2 diabetes

**DOI:** 10.1101/2022.10.11.511721

**Authors:** Joaquín Lilao-Garzón, Yeray Brito-Casillas, Oscar Quesada-Canales, Ana M Wägner, Silvia Muñoz-Descalzo

## Abstract

**Aims/hypothesis:** Delayed maternal age, obesity and diabetes are associated with reduced fertility. We investigated how age and obesity/metabolic syndrome impact fertility and hypothesized that its decrease is due to defects in preimplantation embryo development.

**Methods:** Three groups of female C57Bl6 mice (12 weeks, 9 months and 1 year old) were fed either a high fat diet for 8 weeks, to induce obesity and the metabolic syndrome, or a control chow diet. Body weight and composition, glucose tolerance and insulin resistance were assessed. Fecundity was evaluated by mating and pregnancy rates, as well as number of embryos. Embryo quality was assessed morphologically, and cell fate composition was analysed in preimplantation embryos by state-of-the-art single cell quantitative confocal image analysis.

**Results:** The high fat diet was associated with increased adiposity, glucose intolerance and insulin resistance, especially in the older mice. Fecundity was affected by age, more than by the diet. Both age and high fat diet were associated with reduced cell fate allocation, indicating a delay in preimplantation embryo development, and with increased expression of GATA3, an inhibitor of placentation.

**Conclusion/Interpretation:** These results support that age and the metabolic syndrome reduce fertility through mechanisms which are present at conception very early in pregnancy.

**What is already known about this subject?:**

- Lifestyle changes in modern societies have led to an increase in obesity and type 2 diabetes, and women tend to become pregnant later than ever. These factors have a negative influence on female fecundity.
- In mice, diet induced obesity is associated with poor quality oocytes that affect overall embryonic development.

**What is the key question?:**

- Do age and high fat diet influence cell fate differentiation during preimplantation embryo development?

**What are the new findings?:**

- Body composition and glucose metabolism are altered due to high fat diet even when weight is not affected in young animals.
- Although there are no differences in mating and fertilization rates, embryo quality is lower with high fat diet.
- Cells not fully committed to a cell fate (epiblast or primitive endoderm) are increased in embryos from mature dams or fed a high fat diet, indicating a delay in preimplantation embryo development.

**How might this impact on clinical practice in the foreseeable future?:**

- Our findings show a delay in early embryo development associated to obesity and maternal age. This delay could be responsible for the low fertility observed in women with type 2 diabetes and obesity.

## Introduction

Lifestyle changes have led to an epidemic increase in obesity and type 2 diabetes, both of which have a negative impact on fertility [1] and offspring health [2]. Intrauterine exposure to diabetes and obesity is associated with short and long-term health risks, such as congenital anomalies and stillbirth, neurodevelopmental disorders, obesity, heart disease and type 2 diabetes [3–5].

Furthermore, the age at first pregnancy has been delayed in Europe to 29.4 years in 2019 [6] with consequences on reproductive capacity, embryo quality and even offspring’s overall health [7].

Fecundity is defined by the physiological potential to bear offspring while fertility is the actual number of offspring produced [8]. Obesity and glucose dysregulation lead to altered puberty, ovulation and fertility in women [9]. Therefore, women with obesity and/or diabetes have worse outcomes following fertility treatments, usually due to poorer response to gonadotropins and lower yields in harvested oocytes [10]. Moreover, live birth rates can increase up to 9% for every unit decrease in BMI [11].

The aim of this study was to assess the effect of age and obesity/metabolic syndrome on different aspects related to fertility: fecundability, embryo number and quality and embryo cell fate composition. To do this, an established murine model of the metabolic syndrome was used, which underwent further characterisation.

## Material and methods

### Animal Procedures

C57Bl/6J mice were bred in house at the Universidad de Las Palmas de Gran Canaria (ULPGC) at the Institute for Biomedical and Healthcare Research (IUIBS) animal facility. All mice were housed with controlled room temperature (20-24ºC) and relative humidity (55-72%), and a 12-h light-darkness cycle. All animal studies were conducted following National and European regulations (RD 1201/2005, Law 32/2007, EU Directive 2010/63/EU), were approved by the Animal Ethics Committee of the ULPGC and were authorised by the competent authority of the Canary Islands Government (reference number: OEBA-ULPGC_10/2019R1).

On average, female mammals cease to exhibit reproductive cycles by middle age, around 15 months in mouse or 51 years in humans, while reproductive maturity is reached at 42 days in mice and 11.5 years in humans [12]. Female mice were divided into three age groups: young adults at the beginning of their reproductive life (Y:12 weeks old), mature adults (M: 9 months old) and middle-late age adults when they were close to menopause (O: 1 year old). Animals from different cages but with the same date of birth were grouped together and then randomly assigned to one of two diets following ARRIVE guidelines [13]. High fat diet (HFD) in rodents is an established model for the induction of obesity and the metabolic syndrome [14–16]. Half of the animals were fed a standard diet (ND) (Envigo, Global Diet 2014) and the other half a HFD (60% energy from fat, D12492; Research Diets, Brogaarden, Lynge, Denmark) for 8 weeks prior to the target age (Fig. 1). Body weight was measured weekly. At the time of the study, morphometry was performed (head, body and tail length), body mass index (BMI) was calculated [17] and body composition measured, by Time Domain Nuclear Magnetic Resonance using a Minispec lean fat analyzer (Bruker Optics, Inc., The Woodlands, TX). Healthy males of up to 6 months of age, from the same colony, were used for breeding purposes.

**Fig. 1:**
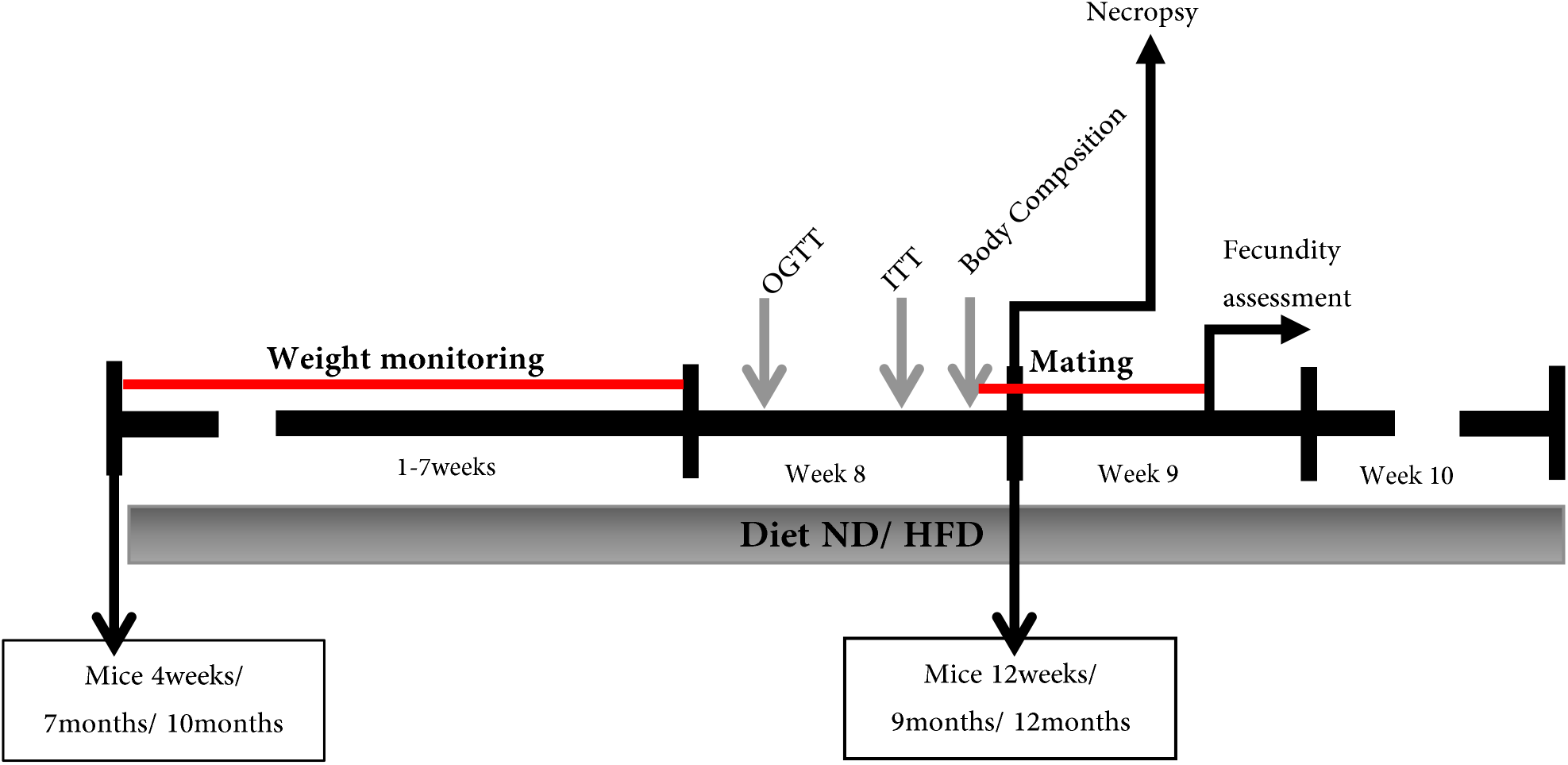
Experiment design. Female mice of different ages were fed either normal diet (ND) or high fat diet (HFD) and their weight was monitored once a week. On the eighth week, oral glucose tolerance test (OGTT), insulin tolerance test (ITT) and body composition assessment were performed. Then, females were mated with young healthy males for up to 8 days and preimplantation embryos were flushed from the uterus 3.5 days after vaginal plugs were detected. A necropsy was performed in a few animals after the 8 weeks of diet.

### Oral Glucose Tolerance Test and Intraperitoneal Insulin Tolerance Test

For the Oral Glucose Tolerance test (OGTT), on the eighth week of diet, mice were fasted for 6h and a 2 g/Kg dose of glucose (G7528, Sigma-Aldrich Chemie GmbH, Steinheim, Germany) was administered by oral gavage. Blood was sampled from the tip of the tail every 15 minutes up to 1 hour and glucose concentrations were measured and recorded (Glucomen Areo, Menarini Diagnostics) [18]. An insulin tolerance test (ITT) was performed by intraperitoneal administration of 0.5 U/Kg of insulin (Actrapid, Novo Nordisk, Copenhagen, Denmark), two days after the OGTT.

### Postmorten Analysis

Animals were killed by isoflurane overdose and bled out from the inferior vena cava. Serum was obtained and a full biochemistry panel comprising albumin, alkaline phosphatase, alanine aminotransferase, amylase, blood urea nitrogen, creatinine, globulin, glucose, phosphorus, total bilirubin, total protein, cholesterol and creatine kinase, was performed (PointCare V2 Biochemistry Analyzer, RAL SA, Barcelona).

Liver, pancreas, heart, spleen, reproductive system, white and brown fat, kidneys, and gastrocnemius muscle were dissected during the necropsy. The tissues were weighed and preserved in 4% neutral buffered formalin (Applichem-Panreac, Barcelona), processed routinely, and embedded in paraffin-wax. Sections (5 μm-thick) were stained with haematoxylin-eosin. All histological sections were randomly and blindly evaluated by a board-certified veterinary pathologist (OQC).

Pancreatic insulin content was evaluated by immunofluorescence. After permeabilization with Triton X-100 0.05%, pancreas sections were incubated with anti-insulin (Santa Cruz, Dallas, TX, USA, sc-9168; 1:80) and anti-Glucagon antibodies (Sigma-Aldrich, G2654, St. Louis, MO, USA, 1:100). Alexa Fluor 488 Tyramide Superboost kit (Invitrogen/Thermo Fisher Scientific, Waltham, MA, USA, B40943) was used to increase insulin fluorescent signal following the manufacturer’s instructions.

### Fecundity Analysis

Animals were mated for up to eight days to allow two consecutive oestrus cycles and pregnancy was confirmed by the presence of a vaginal plug. The ability to mate and get pregnant and the number of embryos per female were used as fecundity indicators. Mating rate was calculated as the proportion of females with a vaginal plug, as an indirect measure of oestrus cycle. Fertilization rate was calculated as the percentage of females with a plug which had embryos in their uteri. Preimplantation embryo quality was assessed by morphological features and classified into 4 categories (A-D) [19].

### Preimplantation embryo analysis

Before implantation, the mammalian embryo develops into a structure known as blastocyst in which three cell types can be identified [20]. The outer cells, the trophectoderm (TE), become the foetal portion of the placenta. The compact inner cells form the inner cell mass (ICM), which will differentiate into Epiblast (Epi) or Primitive Endoderm (PrE). PrE cells generate the endodermal part of the yolk sac while Epi cells form the proper embryo. TE cells are characterized by the expression of markers like GATA3 or CDX2 while in early embryos ICM cells co-express both NANOG and GATA6. During differentiation of the ICM in mid blastocysts, cells asynchronously downregulate one of these markers so that Epi progenitors express NANOG, while PrE progenitors express GATA6 [21] followed by other markers such as SOX17, GATA4, or SOX7 [20]. In late blastocysts, PrE cells migrate to form an epithelial layer of cells next to the blastocoele, leaving the Epi cells in an internal localization. These differentiation processes, occurring prior to implantation, can be assessed using quantitative immunofluorescence analysis (QIF) [22]. After 3.5 days of embryonic development, females were killed and preimplantation embryos were flushed from dissected uteri in M2 medium (Embryomax®; Millipore, Ref. MR-015-D, Burlington, MA, EE. UU) and prepared for immunofluorescence as previously described [23]. Primary antibodies were anti-NANOG (eBioscience, San Diego, CA, USA, Ref.14–5761; 1:200), anti-GATA6 (R&D Systems, Minneapolis, MN, USA, AF1700; 1:200), anti-GATA4 (Santa Cruz, Dallas, TX, USA, sc-9053; 1:200) and anti-GATA3 (eBioscience, San Diego, CA, USA, Ref. MA1-028; 1:200). DAPI (Invitrogen/Thermo Fisher Scientific, Waltham, MA, USA, D1306; 1:1000) was used to stain the nuclei.

### Microscopy and Image Analysis

Both Islets of Langerhans and embryos were imaged using a Zeiss LSM Zeiss LSM700 and a Plan-Apochromat 40x/1.3 Oil DIC (UV) VIS-IR M27 objective, with optical section thickness of 1 μm.

Embryo images were processed and analysed as previously described [22] including the fluorescence decay along the Z axis correction [24]. Briefly, embryos were classified, according to their total cell number, into early (up to 64 cells), mid (65-90 cells) and late (from 91 cells). Cells were then manually classified into TE and ICM. Cells in the ICM were classified using mixture analysis as positive or negative: double negative (DN, negative expression for NANOG and GATA6), Epi progenitors (positive for NANOG and negative for GATA6), PrE progenitors (negative for NANOG and positive for GATA6), and double positive (DP) for uncommitted ICM cells positive for both, NANOG and GATA6. The same classification was based on GATA4 staining instead of GATA6.

### Statistics

Data are presented as mean ± standard error (SEM). Mann-Whitney’s or Student’s t tests were used for comparisons between groups and Z-test for frequencies, respectively, followed by the Bonferroni correction for multiple comparisons. Values of p < 0.05 were considered statistically significant.

## Results

### Eight weeks of HFD are associated with altered body composition and glucose metabolism, especially in older animals

Rapid weight gain was observed in the mature and old groups on HFD, whereas no significant effect on body weight or BMI was seen in young animals (Fig. 2a, b), although HFD was consistently associated with increased total body fat and reduced lean mass (Fig. 2c).

**Fig. 2:**
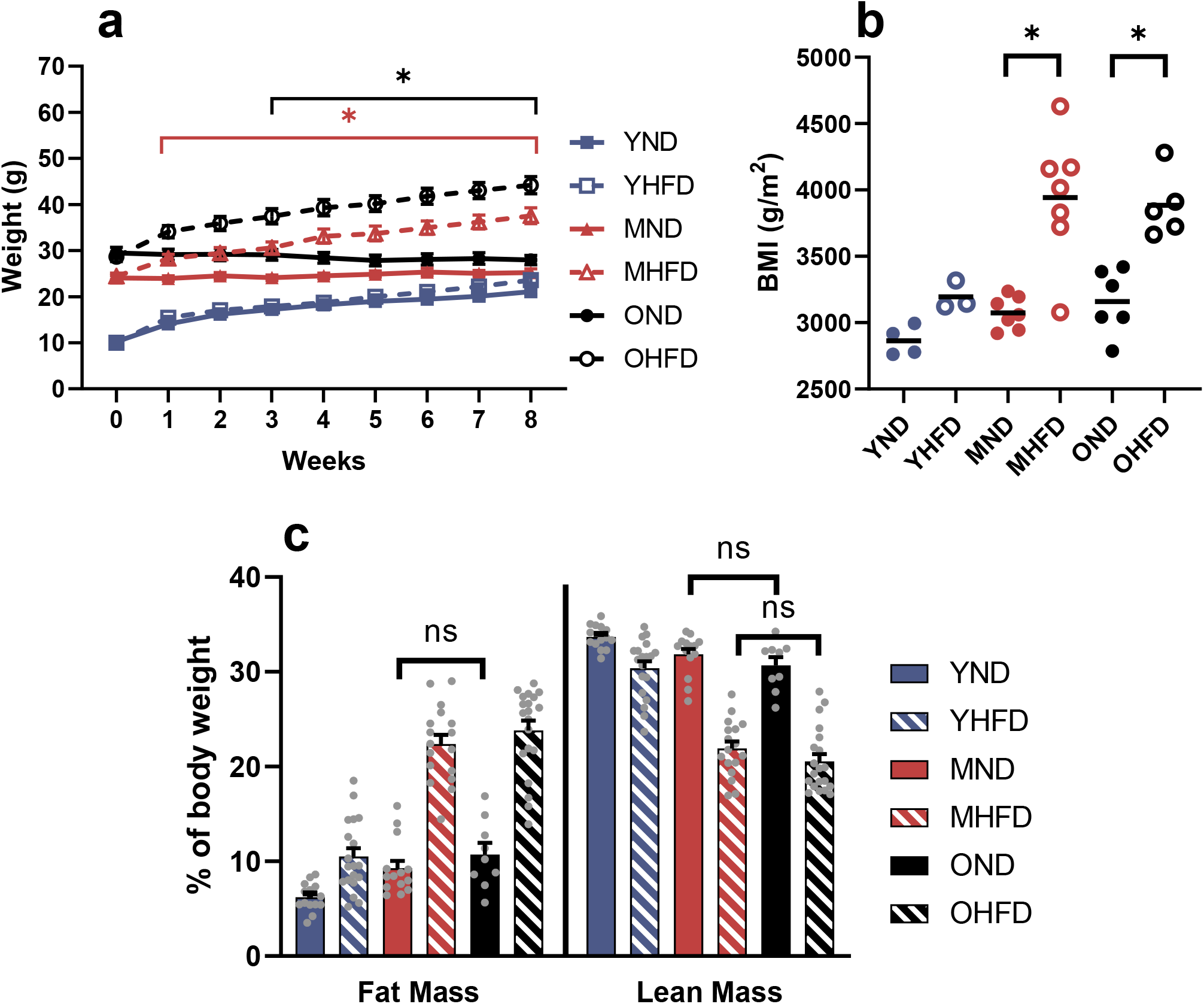
Effect of 8 weeks of high fat diet (HFD) on weight and body composition compared to normal diet (ND). (a) Weekly body weight and (b) body mass index. n= 14 YND, 20 YHFD, 14 MND, 17 MHFD, 9 OND and 21 OHFD. (c) Body composition by nuclear magnetic resonance. n= 14 YND, 20 YHFD, 14 MND, 17 MHFD, 9 OND and 20 OHFD. Data are expressed as mean ± SEM, *p<0.05 comparing each HFD group with its ND control or between age-groups. ns= Not significant, otherwise significant *p<0.05. YND= Young (12 weeks) Normal Diet, YHFD= Young High Fat Diet; MND= Mature (9 months) Normal Diet, MHFD= Mature High Fat Diet; OND= Old (1 year) Normal Diet, OHFD= Old High Fat Diet, here and in subsequent graphs.

OGTTs and ITTs were performed in the last week of diet to evaluate the effect of both age and HFD on glucose metabolism (Fig. 1, 3). The mice fed the HFD showed higher glucose concentrations during the OGTT than those fed ND, and this was especially evident in the older mice (Fig. 3a, b). HFD was also associated with less glucose-lowering after an insulin injection, reflecting increased insulin resistance (Fig. 3c, d).

**Fig. 3:**
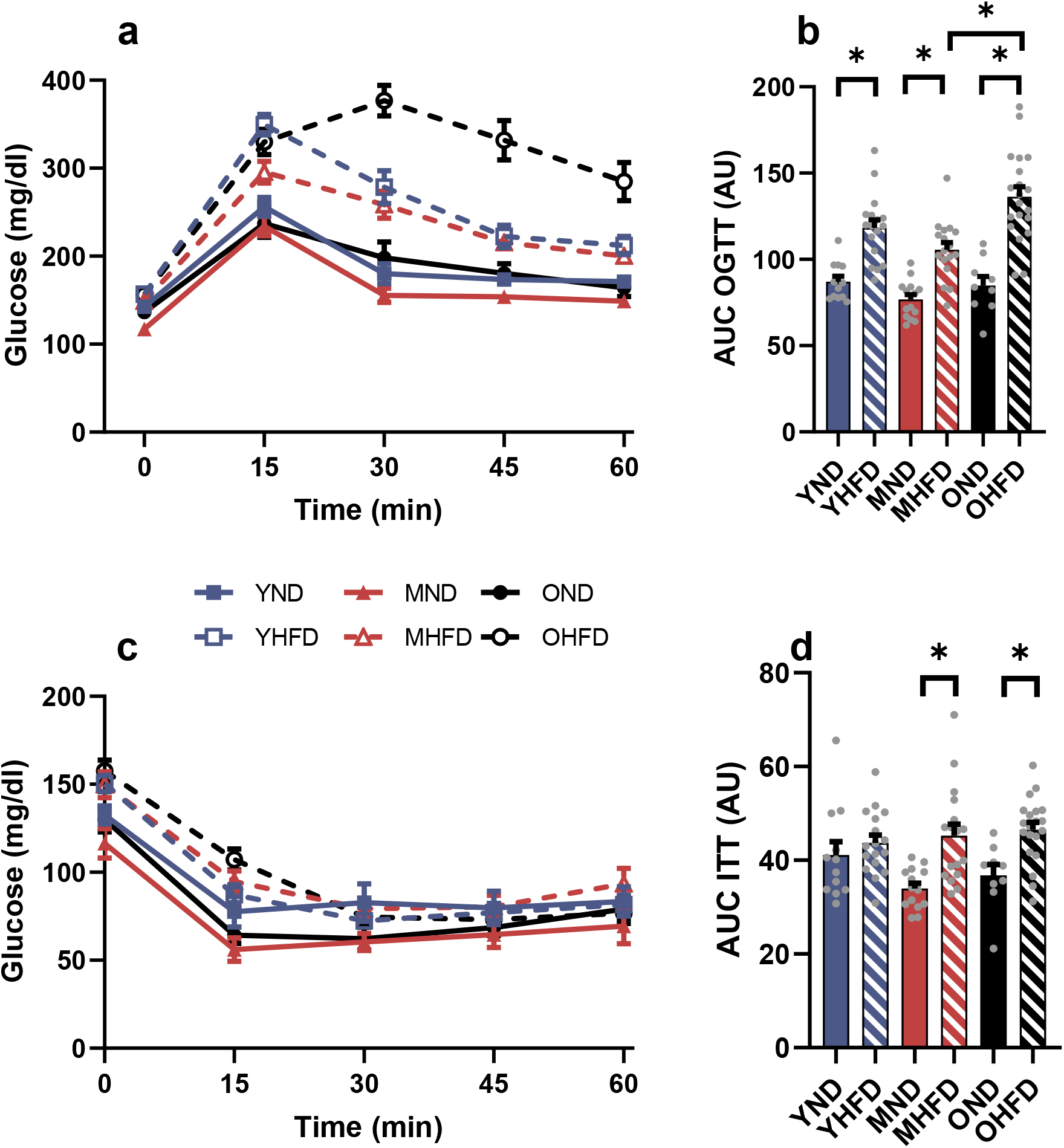
Effect of 8 weeks of high fat diet (HFD) on oral glucose tolerance test (OGTT) and insulin tolerance test (ITT). (a) OGTT blood glucose. (b) AUC calculated from OGTT. (c) ITT blood glucose. (d) AUC calculated from ITT. Data are expressed as mean ± SEM, *p<0.05 comparing each HFD group with its ND control or between age-groups; n=12 YND, 17 YHFD, 14 MND, 17 MHFD, 9 OND and 20 OHFD. YND= Young Normal Diet, YHFD= Young High Fat Diet, MND= Mature Normal Diet, MHFD= Mature High Fat Diet, OND= Old Normal Diet, OHFD= Old High Fat Diet.

### HFD results in increased white fat and reduced muscle mass and high cholesterol and glucose, but no major pathological findings

Post mortem studies confirmed that animals fed the HFD had more body fat and less muscle mass than those fed ND (Fig. 4a, b). A weight increase was also observed in the reproductive system, but this increase was due to the increment of periovaric fat (Sup Fig. 1). Furthermore, high levels of fat in the diet seem to have a weight curbing effect on the liver and pancreas. Regarding blood biochemistry, animals were grouped according to their diet to increase statistical power. Animals fed HFD showed higher glucose and total cholesterol concentrations than those fed ND (Fig. 4c, d). High glucose concentrations were observed even in ND in the blood obtained post-mortem from the inferior vena cava, in agreement with previous reports [25, 26] and is attributed to isofluorane [27]. No differences were observed in other measurements (Sup. Fig. 2).

**Fig. 4:**
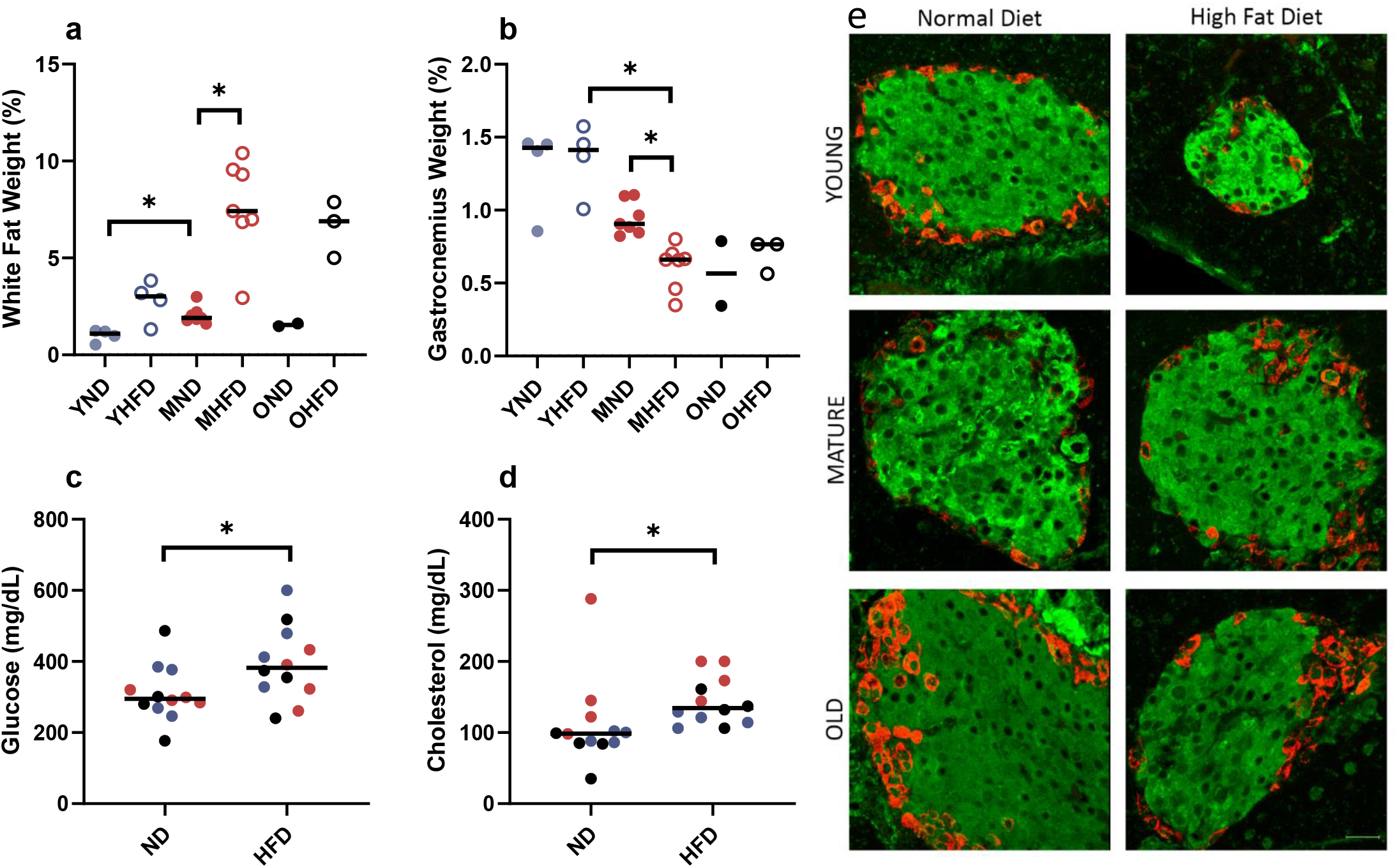
Mice were killed and dissected after being fed for 8 weeks with normal or high-fat diet and their organs were weighed. (a) White fat as the addition of fat from the paragenital fat pads. (b) Gastrocnemius as the addition of the weight of both gastrocnemius muscles from the hind legs. (c-d) For the biochemistry analysis animals were grouped by diet: (c) blood glucose levels and (d) total cholesterol levels at the moment of death. Data are expressed as median, *p<0.05 either comparing each HFD group with its ND control or between ages; Each point refers to one animal. 12 weeks (blue), 9 months (red) and 1 year old (black). (e) Pancreas histological sections immunostained for Insulin (green) and Glucagon (red). Representative maximum projection images from Z stacks for each condition are shown. Scale bar: 20 μm. YND= Young Normal Diet, YHFD= Young High Fat Diet, MND= Mature Normal Diet, MHFD= Mature High Fat Diet, OND= Old Normal Diet, OHFD= Old High Fat Diet.

Histological findings were consistent with senescence and/or the phenotype of the model (Table 1). In absence of an infectious agent, the hepatocellular necrosis observed can be considered as background lesion [28–31]. Also, the systemic amyloidosis observed is considered a common age-related change in some mouse strains, such as the C57BL6 [28] and is consistent with the diagnosis of spontaneous systemic amyloidosis [32].

**Table 1:**
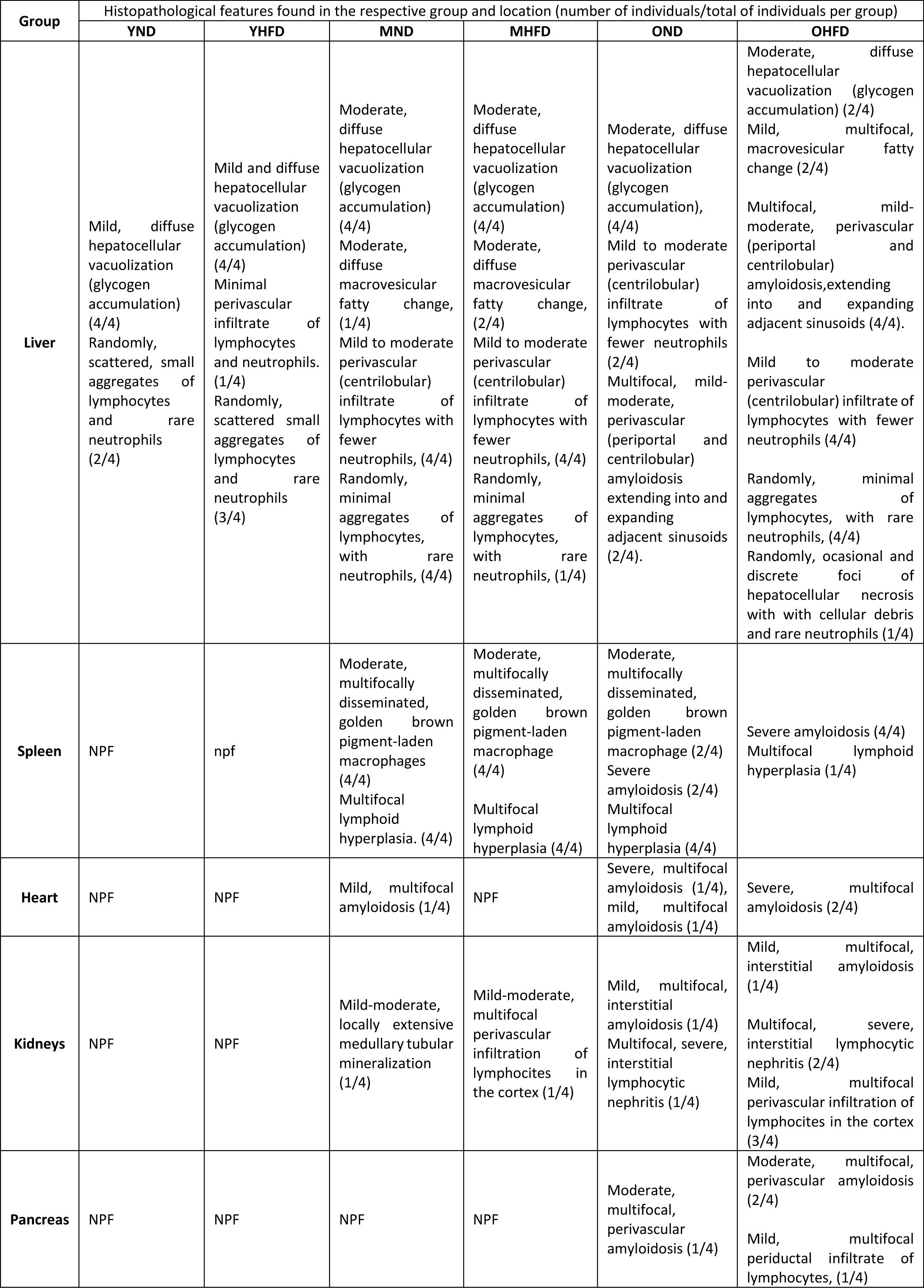
Pathology evaluation of tissue samples. Only tissues with histopathological features are included in this descriptive table; n=24 (4/group). NPF= Non pathological findings; YND= Young Normal Diet, YHFD= Young High Fat Diet, MND= Mature Normal Diet, MHFD= Mature High Fat Diet, OND= Old Normal Diet, OHFD= Old High Fat Diet.

Young groups showed fewer lesions than older groups, but no evident differences were noted between ND and HFD at any age. No qualitative differences were observed in pancreatic insulin or glucagon content comparing age groups or diets (Fig. 4e).

### Overall fecundity decreases with age

Mating and fertilization rates were used as descriptive methods to assess fecundity. Old HFD animals showed lower mating and fertilization rates than younger groups (Fig. 5). Only one old ND dam had embryos and no embryos were obtained in old HFD dams. Hence, the old groups were excluded from further analysis.

**Fig. 5:**
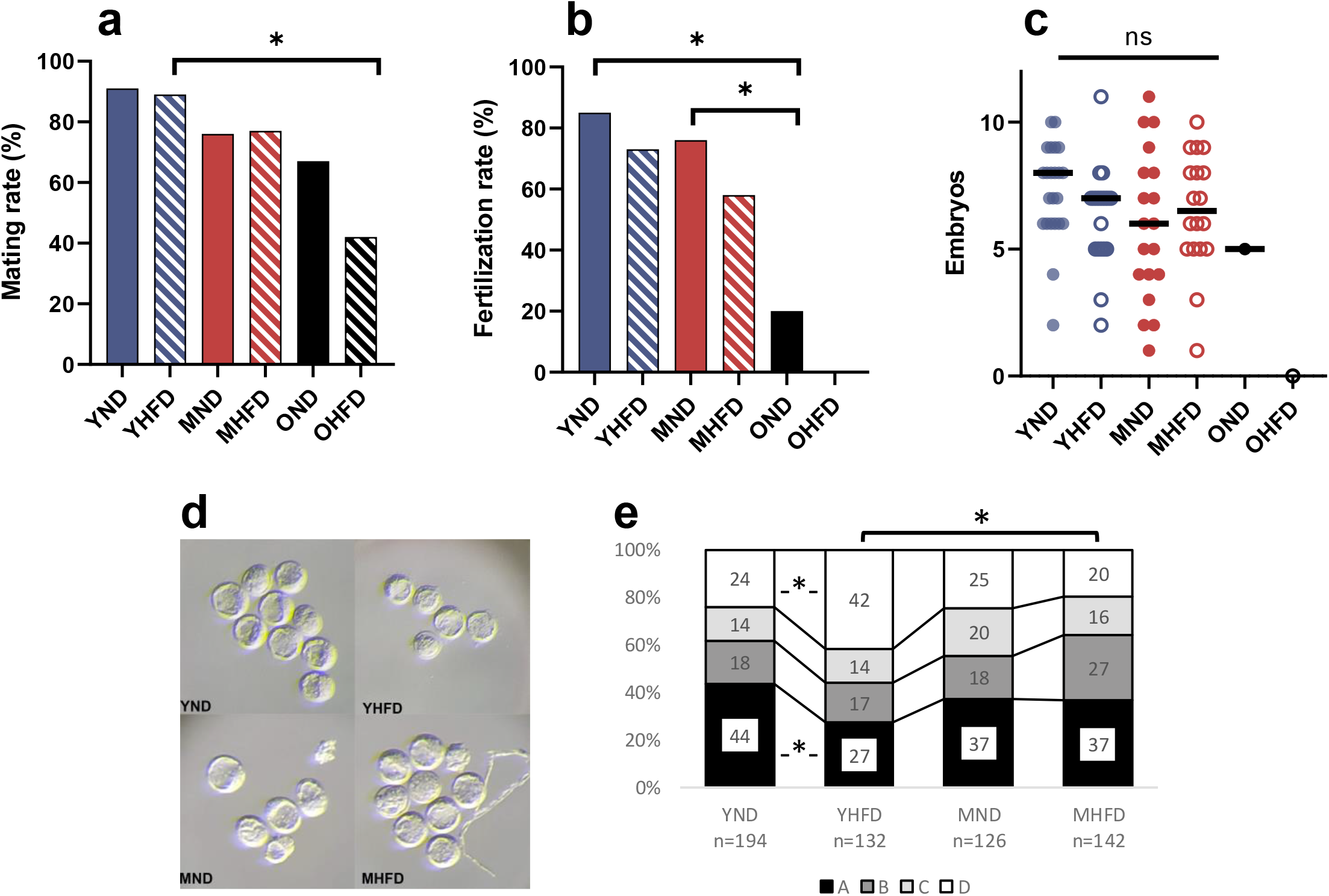
Effect of 8 weeks of high fat diet on fecundity. (a) Mating rate. n= 33 YND, 38 YHFD, 41 MND, 48 MHFD, 9 OND, 20 OHFD. (b) Fertilization rate. n= 27 YND, 26 YHFD, 25 MND, 31 MHFD, 5 OND and 1 OHFD. (c) Number of embryos recorded per female. Each dot indicates one litter. No statistical significance was found in any case, *p<0.05 either comparing each HFD group with its ND control or between ages. (d) Representative micrographs of embryos from indicated females. (e) Embryos classified according to their morphological features into four categories (A, B, C, and D, from high to low quality) *p<0.05; YND= Young Normal Diet, YHFD= Young High Fat Diet, MND= Mature Normal Diet, MHFD= Mature High Fat Diet, OND= Old Normal Diet, OHFD= Old High Fat Diet.

Embryo quality was assessed according to morphological features. Young ND dams had a higher percentage of high quality (category A) embryos and fewer with the worst features (category D) compared with young HFD dams (Fig. 5e). Total cell number together with the classification into TE or ICM cells was done, and the ratio ICM/TE calculated. An age effect was observed in HFD groups (see Sup. Table 1), indicating altered patterns in embryo cell fate allocation.

### Age and HFD are associated with delayed development in preimplantation embryos

Preimplantation embryos flushed from the uteri were stained for GATA3, NANOG and GATA6 and images were analysed by single-cell quantitative methods (Figs. 6-7). Results for early embryos are shown (see classification details in Methods). In general, embryos from HFD females showed an increased proportion of uncommitted cells (DP cells) when compared with their ND controls. This is especially noticeable in mature groups where Epi progenitor proportion diminished upon HFD feeding (Fig. 7a). Comparing embryos from different ages, despite no change in uncommitted cells, mature dams showed a defect in cells to progress towards PrE cell fate.

**Fig. 6:**
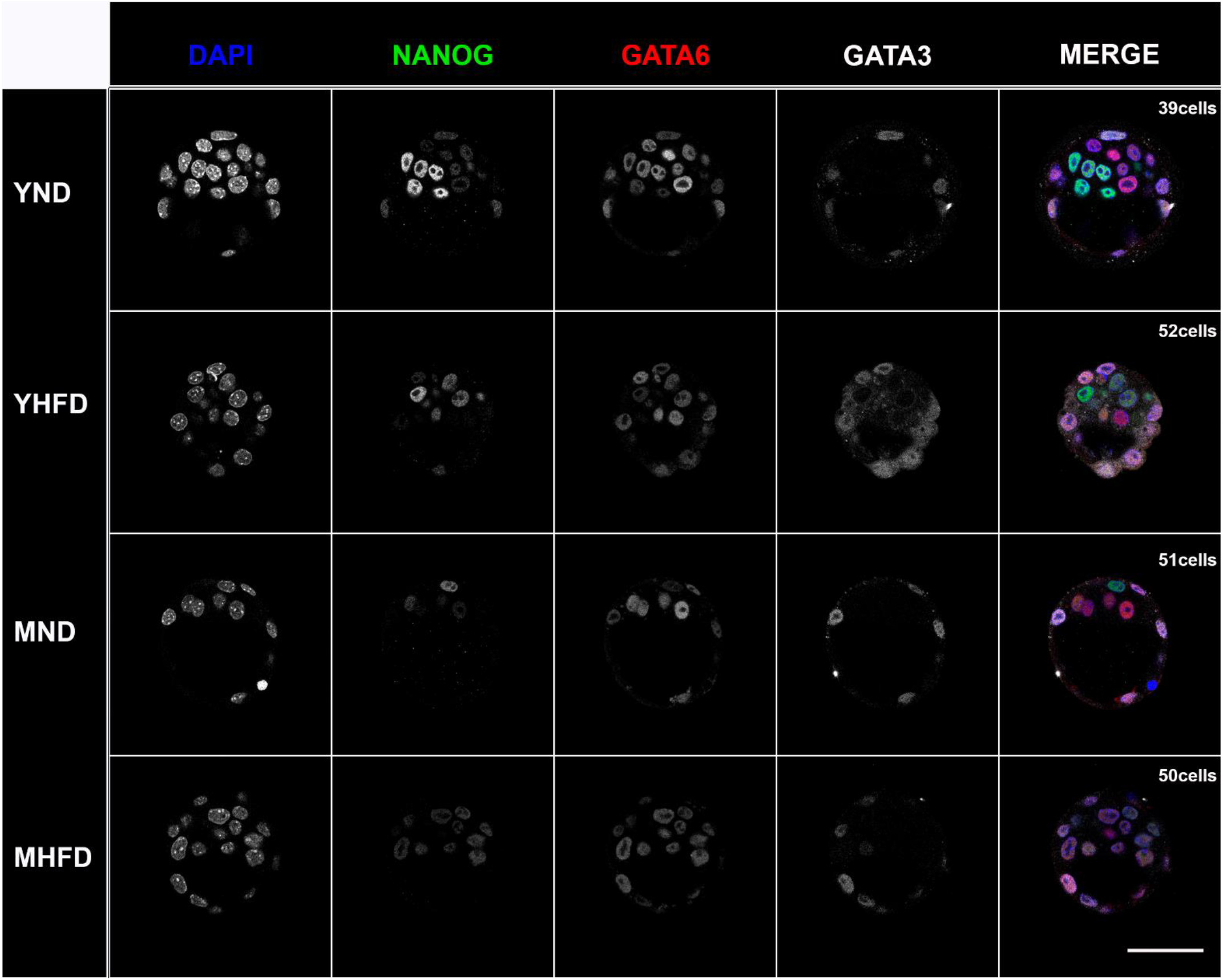
Representative confocal images of mouse preimplantation embryos, immunostained for DAPI (blue), NANOG (green), GATA6 (red) and GATA3 (white) at early stage. Staging criteria: Early (up to 64 cells), Mid (65-90 cells) and Late (from 91 cells). Young Normal Diet (YND); Young High Fat Diet; Mature Normal Diet (MND); and Mature High Fat Diet (MHFD). All embryos shown were immunostained, imaged and processed together. The first four columns are single confocal z-sections; the last column show the merged confocal images. Scale bar: 50 μm.

**Fig. 7:**
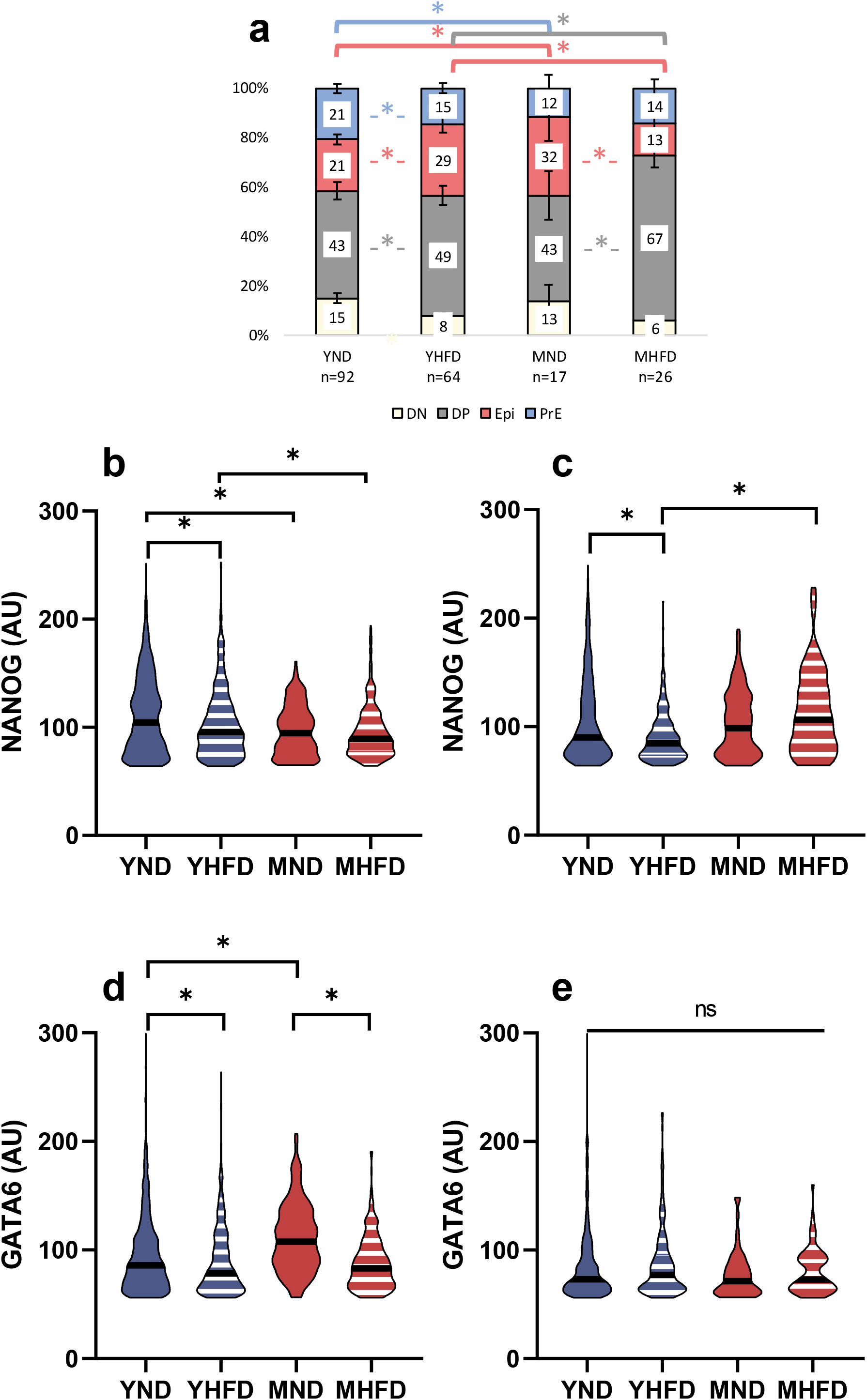
Variations in embryo developmental stage assessed by single cell quantitative immunofluorescence. (a) Population analysis as the percentage of the total number of cells in the ICM. (b-c) Violin plots showing NANOG expression levels in single DP cells (b) and in Epi progenitors (c). (d-e) Violin plots showing GATA6 levels in single DP cells (d) and in PrE progenitors (e). Data are expressed as mean ± SEM, *p<0.05 comparing each HFD group with its ND control or between age-groups. Results showed only for early embryos; YND= Young Normal Diet, YHFD= Young High Fat Diet, MND= Mature Normal Diet, MHFD= Mature High Fat Diet, OND= Old Normal Diet, OHFD= Old High Fat Diet.

Although the population analysis showed more Epi progenitors in young HFD embryos, NANOG levels were significantly lower both in DP cells (Fig. 7b) and in Epi progenitors (Fig. 7c). The same was observed in mature dams regardless of the diet: NANOG levels were significantly lower in DP cells (Fig. 7b). Epi progenitors showed higher levels of NANOG in mature only on HFD (Fig. 7c)

GATA6 levels were significantly lower in HFD embryos independently of the age in DP cells (Fig. 7d). MND embryos showed higher levels of GATA6 than YND embryos (Fig. 7d).

Equivalent results were obtained for the population analysis in mid and late embryos (Sup. Fig. 3) as well as the analysis with the more developmentally advanced PrE marker GATA4 (Sup. Fig. 4).

The higher number of uncommitted cells together with fewer cells fully committed to one cell fate (especially to PrE), and cell fate marker levels indicate that preimplantation embryos developed under an obese environment or from mature females are delayed in their normal development.

### Both HFD and advanced maternal age had an increasing effect in the levels of GATA3

GATA3 was analysed in the TE cells where it was almost ubiquitously expressed (Fig. 8a). GATA3 expression increased with age. However, its response to HFD changed in opposite directions in young (increase) and mature (decrease) dams (Fig. 8b).

**Fig. 8:**
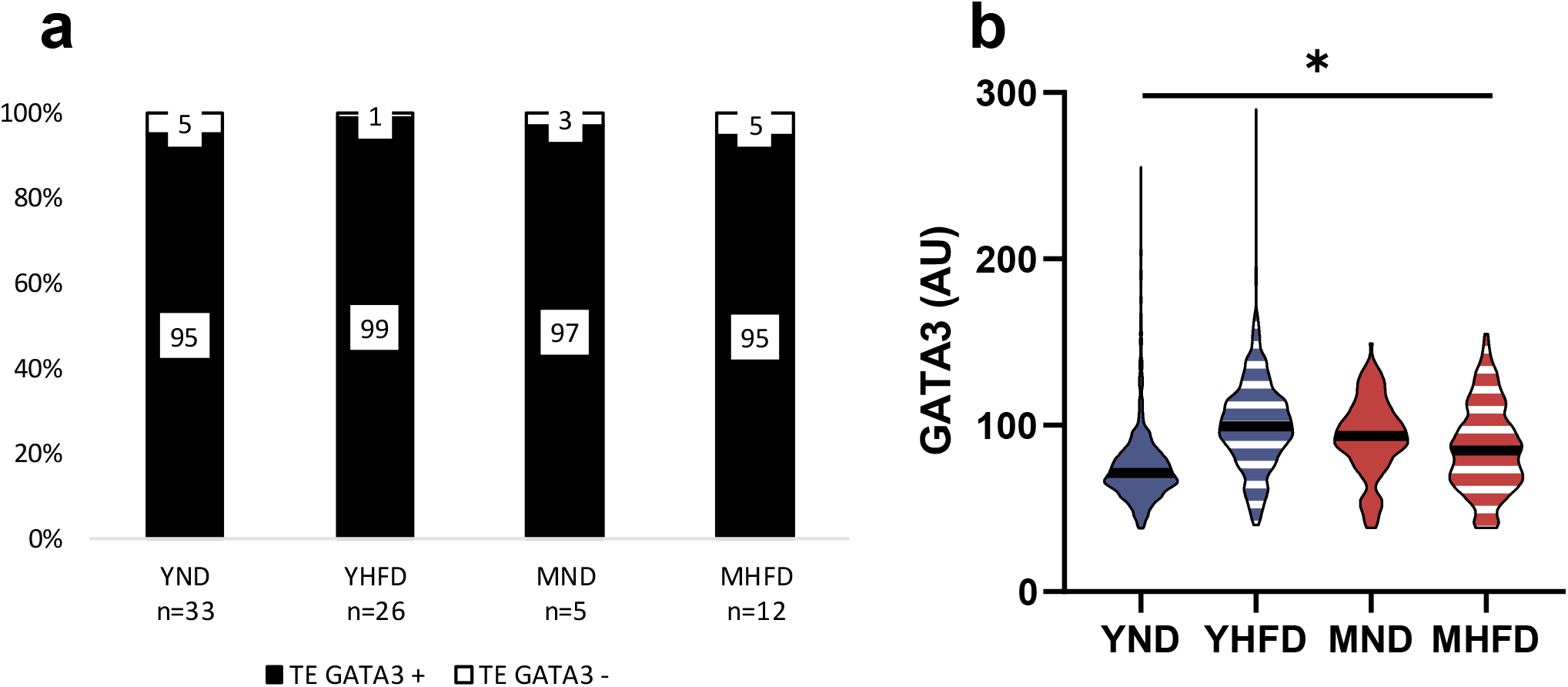
Variations in GATA3 levels in the trophectoderm (TE) assessed by quantitative immunofluorescence. (a) Population analysis as the percentage of the total number of cells in the TE. (b) Violin plots showing GATA3 expression levels. *p<0.05 either comparing each HFD group with its ND control or between ages. Results showed only for early embryos; YND= Young Normal Diet, YHFD= Young High Fat Diet, MND= Mature Normal Diet, MHFD= Mature High Fat Diet.

## Discussion

In the present study, we assessed the impact of maternal age and HFD on early embryo development, specifically Epi versus PrE cell fate decision, through state-of-the-art single cell quantitative confocal image analysis. Indeed, HFD led to increased body fat, glucose intolerance and insulin resistance. To our knowledge, this is the first study to report delayed early embryo development, based on Epi versus PrE cell fate specification, associated with both HFD and age. We also describe alterations in GATA3 levels, showing an effect of age and HFD in TE differentiation. This is significant as it allows us to establish embryo development disorders that could be responsible for embryo loss in the first few days after fertilization, leading to the low fertility described in older and overweight women [10].

We first confirmed that 8 weeks of HFD in adulthood induced overweight in female mice as shown in previous studies [33, 34] but not when started before puberty. However, despite similar weight and BMI, the young HFD animals showed almost twice as much fat mass as their control group. The OGTT and ITT revealed that HFD induces glucose impairment and insulin resistance, especially in the older groups. However, age itself did not have a direct effect on glucose intolerance, in agreement with a previous study [34].

Frequently, when the effects of childhood obesity in adulthood are assessed, only body weight and BMI are used as indicators [35]. Here, we show that in young mice, 8 weeks of HFD did not affect body weight and BMI. However, other parameters that affect overall health such as body composition and glucose metabolism were severely affected.

Furthermore, we validated the 8 weeks HFD in C57Bl6 female mice as a prediabetic mouse model with different levels of obesity and hyperglycaemia depending on age; young adults, mature adults, and middle-late age adults. This strategy allowed us to evaluate how HFD and maternal age affect fecundity.

Selected maternal ages were based on previous studies. Mice are often divided into mature adults (3-6 months), middle aged (10-15 months) and old (18-24 months) [36, 37] with some authors even expanding mouse lifespan up to 36 months [38]. In this context, their reproductive senescence is established between 10 to 15 months. Other authors establish the irregular fertility that precedes menopause at approximately 8 months of age [39] in accordance with the lower fertility observed in mature groups in the present study.

Infertility (failure to conceive after attempting at least twelve months of natural fertilization) is a rising problem in our society. Life factors such as psychological stress, smoking, drugs or diet and variations in body weight have a substantial effect. Overweight in women is often associated with multiple reproductive disorders such as polycystic ovary syndrome, infertility, increased miscarriage rate, as well as pregnancy complications (gestational diabetes, pre-eclampsia, and macrosomia). Furthermore, overweight (BMI>25 Kg/m^2^) and obese (BMI>30 Kg/m^2^) women have worse outcomes following fertility treatments than women with normal BMI. They respond poorly to induction of ovulation, require higher doses of gonadotropins and longer treatment courses for follicular development and ovulatory cycles [40]. When focusing on their blastocyst quality, high BMI is not associated with low embryo quality, though implantation rate is reduced [41–43]. Type 2 diabetes is most frequently associated with obesity. However, in the absence of obesity, type 2 diabetes is also associated with reduced fertility and more frequent need for assisted reproduction [44]. Implantation rate is indeed reduced in women with type 2 diabetes, regardless of BMI, as shown by a recent study based on the Danish registry of assisted reproductive techniques [43]. Our results showed no significant differences in mating or fertilization rates. However, we did observe reduced embryo quality in YHFD females. Surprisingly, no diet effect was observed in embryos from mature females suggesting that age itself has a bigger impact than obesity.

During mammalian preimplantation development, two sequential cell fate decisions occur. The first decision induces TE (precursor of the placenta) versus ICM cells. The ICM cells make a further decision, differentiating either into Epi or into PrE [20]. Evidence suggests that inflammation and oxidative stress can severely affect these cellular events and embryo quality [45]. Moreover, preimplantation embryos cultivated with lipopolysaccharides showed a lower proportion of SOX17 expressing cells, suggesting that the oxidative stress induced by lipopolysaccharides could be impairing the normal development of the embryonic PrE. Furthermore, the correction of the associated oxidative stress, reverted these PrE development impairments [45].

GATA3 is highly expressed in the invasive murine trophoblast giant cells of the blastocyst and its knockdown enhances placental cell invasion [46]. Hence, the high levels of GATA3 associated with HFD we observe here might impair placentation. Similarly, embryos treated with lipopolysaccharides also showed an increased expression of TE markers [45].

A limitation of this study is that we did not measure caspase-3 activity or reactive oxygen species (ROS) in the embryos to assess oxidative stress. However, previous studies showed that oxidative stress is strongly correlated with the prevalence of type 2 diabetes, obesity and aging through different mechanisms such as high ROS production caused by hyperglycaemia and oxidation of fatty acids or by the decrease in antioxidant capacity [47–50].

In summary, we demonstrated that 8 weeks of HFD are enough to model obesity and the metabolic syndrome in mice even at young ages when there are no apparent effects on weight. Age seems to be the most important factor determining reduced fecundity, but both HFD and age are associated with delayed preimplantation embryo development that might explain the lower implantation rates observed in women with overweight or type 2 diabetes [41, 43].

## Supplementary figures/tables

**Sup. Fig. 1:**
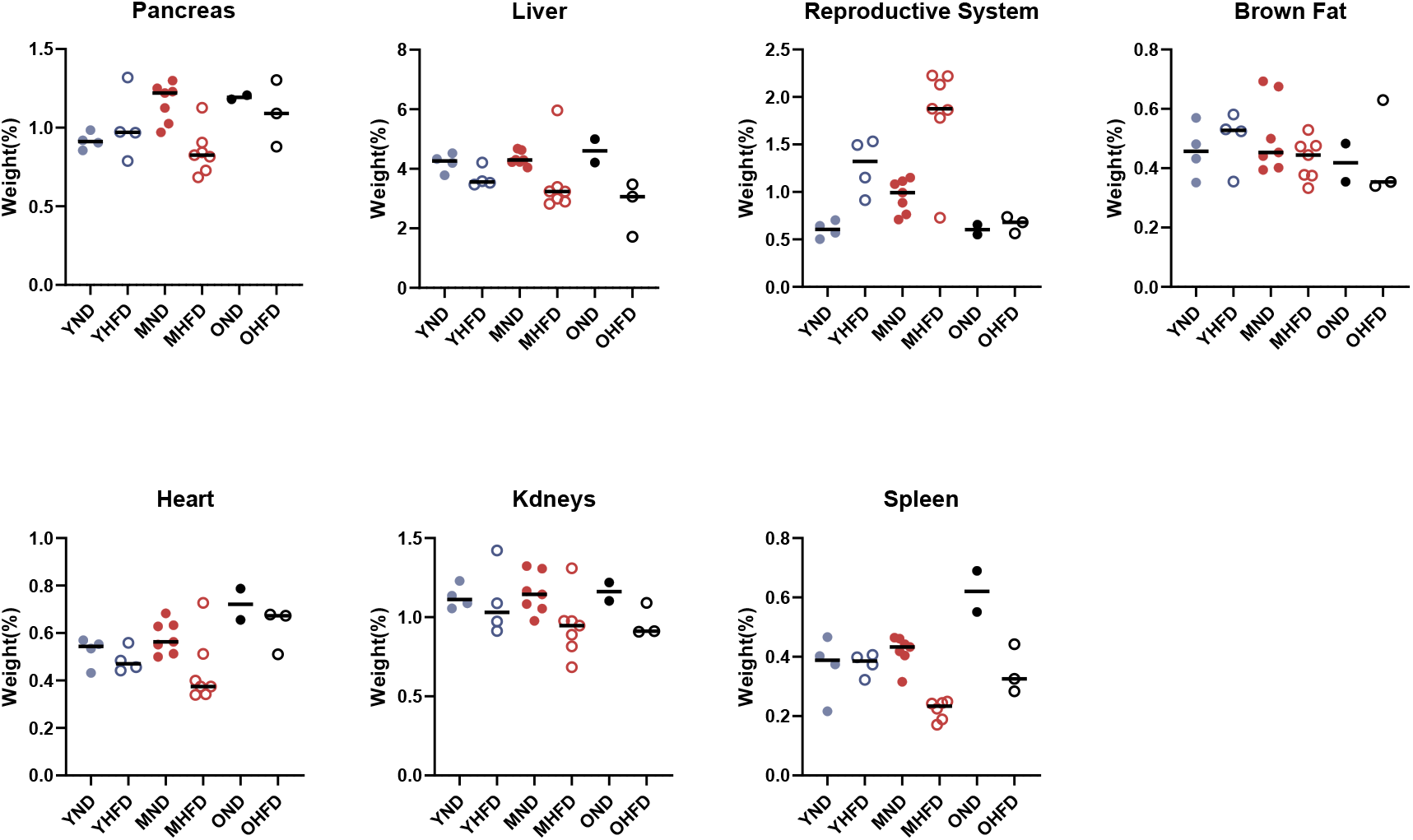
Mice were killed and dissected after being fed for 8 weeks with HFD and their organs were weighed. Pancreas, Liver, Reproductive system, Brown fat, Heart, Kidneys and Spleen; Each point refers to one animal; YND= Young Normal Diet, YHFD= Young High Fat Diet, MND= Mature Normal Diet, MHFD= Mature High Fat Diet, OND= Old Normal Diet, OHFD= Old High Fat Diet.

**Sup. Fig. 2:**
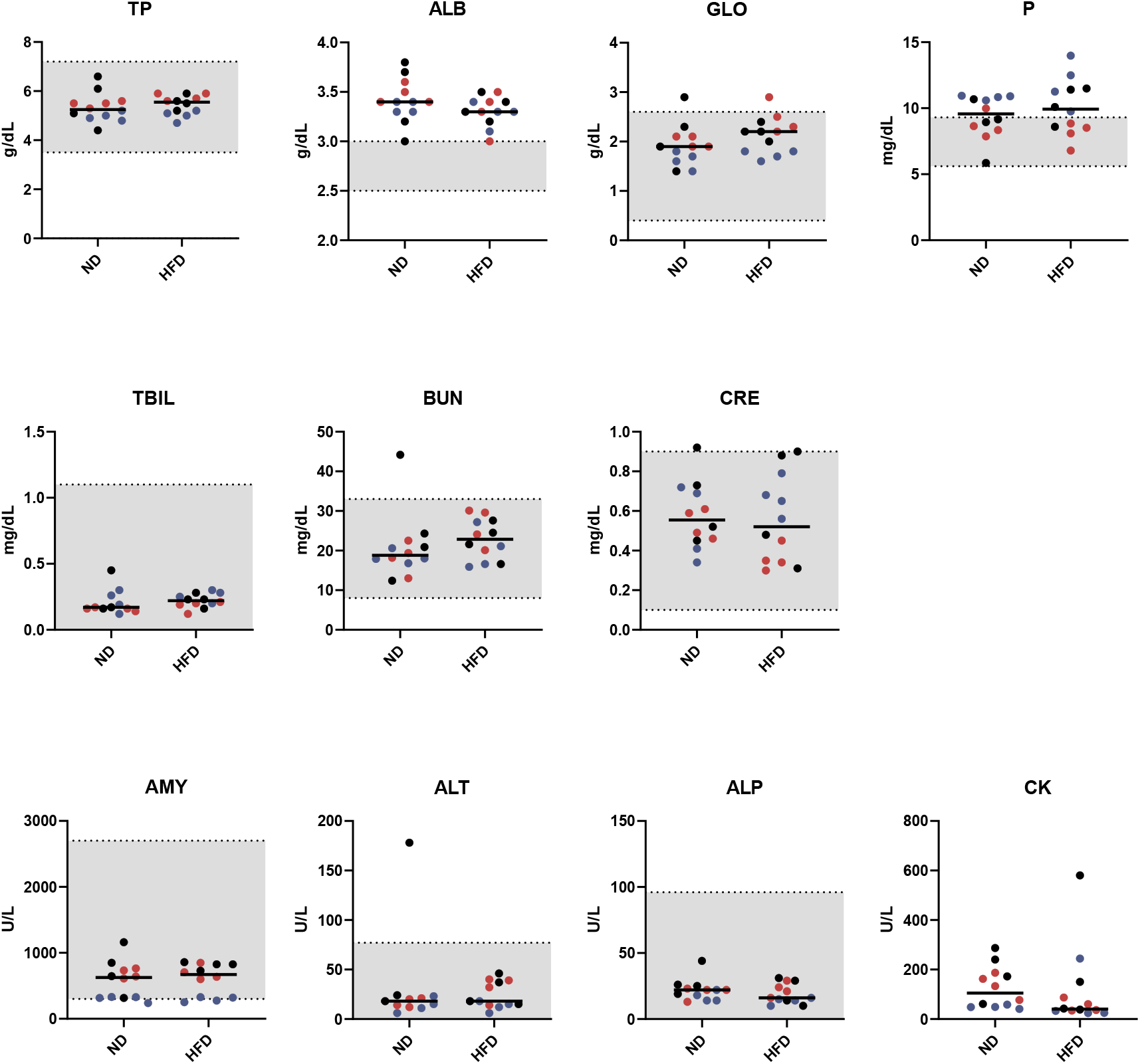
Biochemistry analysis. Animals were grouped by diet: total protein (TP), albumin (ALB), globulin (GLOB), phosphorus (P), total bilirubin (TBIL), blood urea nitrogen (BUN), creatinine (CRE), amylase (AMY), alanine aminotransferase (ALT), alkaline phosphatase (ALP), and creatine kinase (CK). Data are expressed as median, *p<0.05 comparing each HFD group with its ND control; Each point refers to one animal; 12 weeks (blue), 9 months (red) and 1 year old (black). YND= Young Normal Diet, YHFD= Young High Fat Diet, MND= Mature Normal Diet, MHFD= Mature High Fat Diet, OND= Old Normal Diet, OHFD= Old High Fat Diet. Gray areas indicate physiological ranges according to the manufacturer.

**Sup. Table 1:** Embryo analysis by quantitative immunofluorescence. Staging criteria: Early (up to 64 cells), Mid (65-90 cells) and Late (from 91 cells). YND= Young Normal Diet, YHFD= Young High Fat Diet, MND= Mature Normal Diet, MHFD= Mature High Fat Diet. TE= trophectoderm cells, ICM= inner cell mass cells.

|  |  |  | Total cell number |  | Total TE number |  | Total ICM |  | ICM/TE Ratio |  |
| --- | --- | --- | --- | --- | --- | --- | --- | --- | --- | --- |
|  | Stage | N Embryos | Average | SE | Average | SE | Average | SE | Average | SE |
| YND | Early | 92 | 54.18 | 1.06 | 33.93 | 0.71 | 20.25 | 0.56 | 0.61 | 0.02 |
|  | Mid | 33 | 72.12 | 1.07 | 48.82 | 1.23 | 23.30 | 0.68 | 0.49 | 0.03 |
|  | Late | 2 | 92.00 | 1.00 | 68.50 | 3.50 | 23.50 | 2.50 | 0.35 | 0.06 |
| YHFD | Early | 64 | 52.47 | 1.39 | 32.45 | 1.02 | 20.02 | 0.61 | 0.64* | 0.02 |
|  | Mid | 7 | 71.57 | 2.22 | 45.86 | 3.26 | 25.71 | 1.60 | 0.59 | 0.07 |
|  | Late | 1 | 91.00 | NA | 65.00 | NA | 26.00 | NA | 0.40 | NA |
| MND | Early | 17 | 45.35 | 3.39 | 28.47 | 2.19 | 16.88 | 1.51 | 0.60 | 0.04 |
|  | Mid | 4 | 79.00 | 3.54 | 53.00 | 4.02 | 26.00 | 1.08 | 0.50 | 0.06 |
|  | Late | 2 | 93.00 | 0.00 | 67.50 | 0.50 | 25.50 | 0.50 | 0.38 | 0.01 |
| MHFD | Early | 26 | 44.84 | 2.61 | 29.24 | 1.63 | 15.60 | 1.12 | 0.53* | 0.03 |
|  | Mid | 23 | 75.09 | 1.91 | 48.04 | 1.82 | 27.04 | 0.76 | 0.58 | 0.03 |
|  | Late | 14 | 97.86 | 1.62 | 69.21 | 1.97 | 28.57 | 1.45 | 0.42 | 0.03 |

**Sup. Fig. 3:**
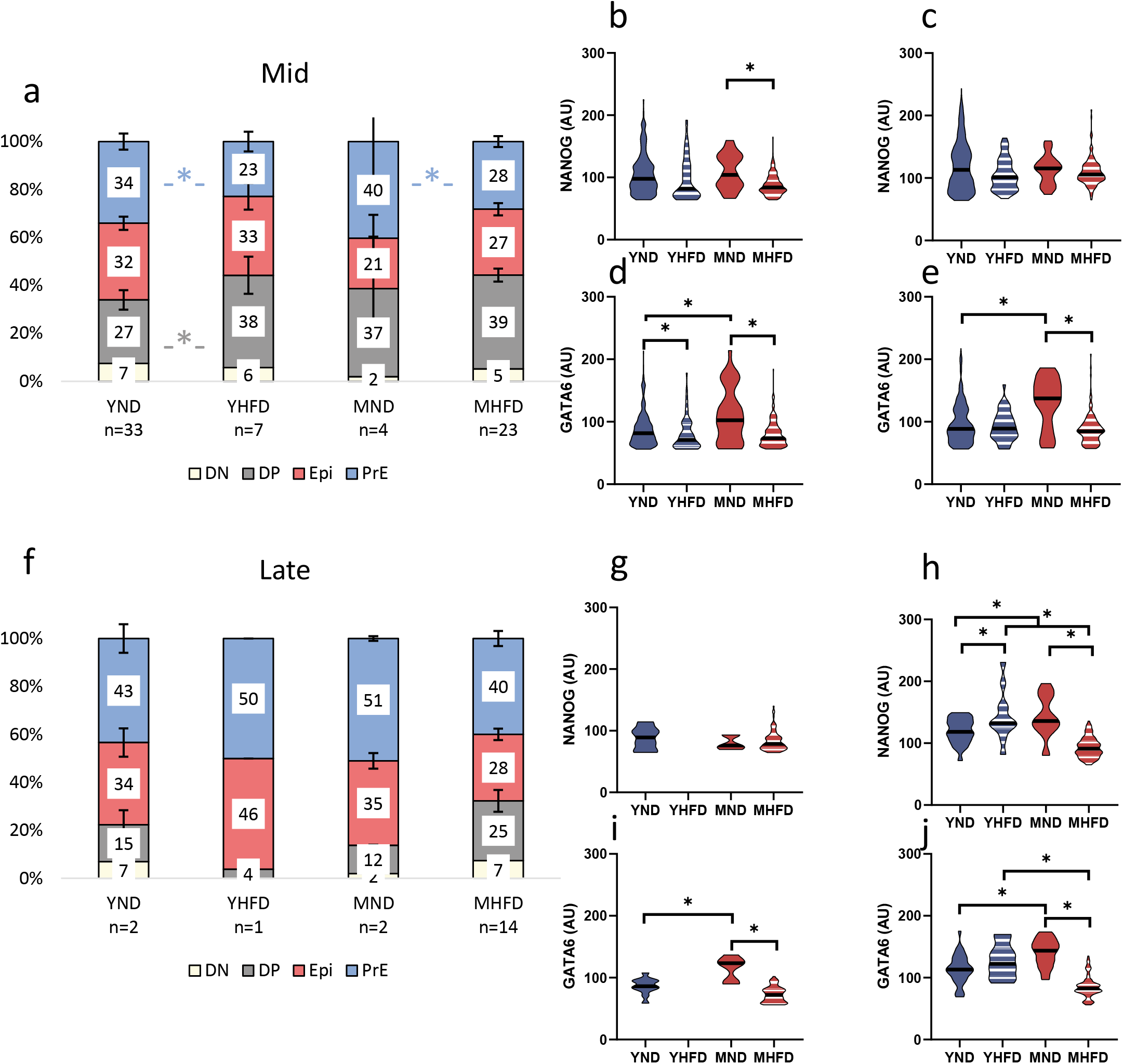
Variations in embryo developmental stage assessed by quantitative immunofluorescence. (a, f) Population analysis as the percentage of the total number of cells in the ICM for Mid (a) and late (f) embryos; Violin plots showing fate marker expression levels: (b, g) NANOG levels in single DP cells and (c, h) in Epi progenitors. (d, i) GATA6 levels in single DP cells (e, j) and in PrE progenitors. Data are expressed as mean ± SEM, *p<0.05 comparing each HFD group with its ND control or between age-groups; YND= Young Normal Diet, YHFD= Young High Fat Diet, MND= Mature Normal Diet, MHFD= Mature High Fat Diet, OND= Old Normal Diet, OHFD= Old High Fat Diet.

**Sup. Fig. 4:**
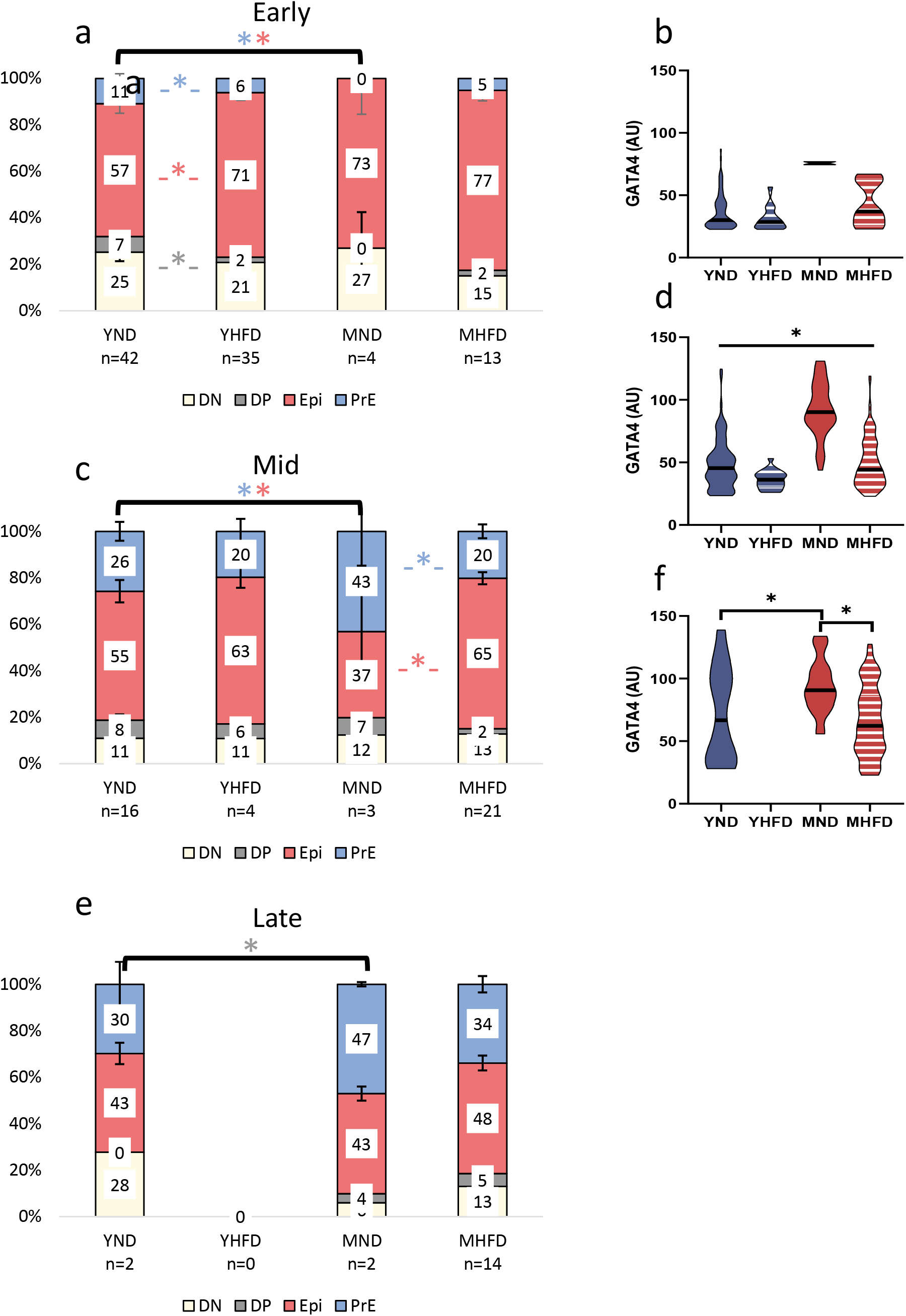
Variations in embryo developmental stage assessed by quantitative immunofluorescence. (a, c, e) Population analysis as the percentage of the total number of cells in the ICM for early (a), Mid (c) and Late (e) embryos; Cells positive for GATA4 but negative for NANOG were considered PrE progenitors. Violin plots showing fate marker expression levels: (b, d, e) GATA4 levels in single PrE progenitors. Data are expressed as mean ± SEM, *p<0.05 comparing each HFD group with its ND control or between age-groups; YND= Young Normal Diet, YHFD= Young High Fat Diet, MND= Mature Normal Diet, MHFD= Mature High Fat Diet, OND= Old Normal Diet, OHFD= Old High Fat Diet.

## Abbreviations

BMI: Body Mass Index
DN: Double Negative
DP: Double Positive
Epi: Epiblast
HFD: High Fat Diet
ICM: Inner Cell Mass
ITT: Insulin Tolerance Test
MHFD: Mature High Fat Diet
MND: Mature Normal Diet
ND: Normal Chow Diet
OGTT: Oral Glucose Tolerance Test
OHFD: Old High Fat Diet
OND: Old Normal Diet
OQC: Oscar Quesada-Canales
PrE: Primitive Endoderm
QIF: Quantitative Immunofluorescence Analysis
ROS: Reactive Oxygen Species
TE: Trophectoderm
YHFD: Young High Fat Diet
YND: Young Normal Diet

## Contribution statement

JLG, YBC, AMW and SMD conceived and designed the research; JLG, YBC, OQC and SMD performed the research, acquired the data and analysed the results; JLG, YBC, SMD and AMW analysed and interpreted the data. All authors were involved in drafting and revising the manuscript.

## Acknowledgements

We are grateful to Tina Balayo, Ana Exposito-Montesdeoca and the IUIBS animal facility staff for the technical support. Sabine Fischer for data analysis support. JLG is supported by the ULPGC predoctoral program. SMD by the “Viera y Clavijo” Program from the Agencia Canaria de Investigación, Innovación y Sociedad de la Información (ACIISI) and the ULPGC. Research at the SMD lab is supported by the ACIISI (CEI2019-02), Programa de Ayudas a la Investigación de la ULPGC, and ACIISI co-funded by FEDER Funds (ProID2020010013). AMW receives research funds from Instituto de Salud Carlos III (PI16/00587, PI20/00846, PMP21/00069) and ACIISI (ProID2021010143) co-funded by European Regional Development Funds, as well as the European Horizon 2020 Programme (101017385) and Fundación Canaria del Instituto de Investigaciones Sanitarias de Canarias (PI19/30, PIFIISC20/16).

## References

1. Thong EP, Codner E, Laven JSE, Teede H (2020) Diabetes: a metabolic and reproductive disorder in women. Lancet Diabetes Endocrinol 8(2):134–149. https://doi.org/10.1016/S2213-8587(19)30345-6

2. Fleming TP, Watkins AJ, Velazquez MA, et al (2018) Origins of lifetime health around the time of conception: causes and consequences. Lancet (London, England) 391(10132):1842. https://doi.org/10.1016/S0140-6736(18)30312-X

3. Faruque S, Tong J, Lacmanovic V, Agbonghae C, Minaya DM, Czaja K (2019) The Dose Makes the Poison: Sugar and Obesity in the United States – a Review. Polish J Food Nutr Sci 69(3):219

4. Godfrey KM, Reynolds RM, Prescott SL, et al (2017) Influence of maternal obesity on the long-term health of offspring. Lancet Diabetes Endocrinol. 5:53–64

5. Ornoy A, Reece EA, Pavlinkova G, Kappen C, Miller RK (2015) Effect of maternal diabetes on the embryo, fetus, and children: Congenital anomalies, genetic and epigenetic changes and developmental outcomes. Birth Defects Res Part C - Embryo Today Rev 105(1):53–72. https://doi.org/10.1002/bdrc.21090

6. Women in the EU are having their first child later - Products Eurostat News - Eurostat. https://ec.europa.eu/eurostat/web/products-eurostat-news/-/ddn-20210224-1. Accessed 24 Mar 2022

7. Myrskylä M, Fenelon A (2012) Maternal Age and Offspring Adult Health: Evidence From the Health and Retirement Study. Demography 49(4):1231–1257. https://doi.org/10.1007/s13524-012-0132-x

8. Shenk M (2015) Fertility and fecundity. Int. Encycl. Hum. Sex. 369–426

9. Nandi A, Poretsky L (2013) Diabetes and the female reproductive system. Endocrinol Metab Clin North Am 42(4):915–946. https://doi.org/10.1016/J.ECL.2013.07.007

10. Silvestris E, de Pergola G, Rosania R, Loverro G (2018) Obesity as disruptor of the female fertility. 16(1):1–13

11. Grzegorczyk-Martin V, Fréour T, de Bantel Finet A, et al (2020) IVF outcomes in patients with a history of bariatric surgery: a multicenter retrospective cohort study. Hum Reprod 35(12):2755–2762. https://doi.org/10.1093/HUMREP/DEAA208

12. Men and mice: Relating their ages | Elsevier Enhanced Reader. https://reader.elsevier.com/reader/sd/pii/S0024320515300527?token=CE03E74B95ABE77B63A2F62BCA8378DF86349C4B6FF3AAADDAB8F3DD3A6D7D87DE97AAB1B3DFCC2ECD1A6157C30F3BF7&originRegion=eu-west-1&originCreation=20220216110724. Accessed 16 Feb 2022

13. Percie Du Sert N, Hurst V, Ahluwalia A, et al (2020) The ARRIVE guidelines 2.0: Updated guidelines for reporting animal research. BMC Vet Res 16(1):1–7. https://doi.org/10.1186/S12917-020-02451-Y/TABLES/2

14. Chen T, Hill JT, Moore TM, et al (2020) Lack of skeletal muscle liver kinase B1 alters gene expression, mitochondrial content, inflammation and oxidative stress without affecting high-fat diet-induced obesity or insulin resistance. Biochim Biophys Acta - Mol Basis Dis 165805. https://doi.org/10.1016/j.bbadis.2020.165805

15. Igosheva N, Abramov AY, Poston L, et al (2010) Maternal Diet-Induced Obesity Alters Mitochondrial Activity and Redox Status in Mouse Oocytes and Zygotes. PLoS One 5(4):e10074–e10074. https://doi.org/10.1371/journal.pone.0010074

16. Lilao-Garzón J, Valverde-Tercedor C, Muñoz-Descalzo S, Brito-Casillas Y, Wägner AM (2020) In Vivo and In Vitro Models of Diabetes: A Focus on Pregnancy. In: Advances in experimental medicine and biology. Adv Exp Med Biol

17. Gargiulo S, Gramanzini M, Megna R, et al (2014) Evaluation of growth patterns and body composition in c57bl/6j mice using dual energy x-ray absorptiometry. Biomed Res Int 2014. https://doi.org/10.1155/2014/253067

18. Brito-Casillas Y, Melián C, Wägner AM (2016) Study of the pathogenesis and treatment of diabetes mellitus through animal models. Endocrinol y Nutr 63(7):345–353. https://doi.org/10.1016/j.endonu.2016.03.011

19. Gardner DK, Balaban B (2016) Assessment of human embryo development using morphological criteria in an era of time-lapse, algorithms and “OMICS”: is looking good still important? Mol Hum Reprod 22(10):704–718. https://doi.org/10.1093/MOLEHR/GAW057

20. Rossant J (2018) Genetic Control of Early Cell Lineages in the Mammalian Embryo. https://doi.org/101146/annurev-genet-120116-024544 52:185–201. https://doi.org/10.1146/ANNUREV-GENET-120116-024544

21. Saiz N, Williams KM, Seshan VE, Hadjantonakis A-K (2016) Asynchronous fate decisions by single cells collectively ensure consistent lineage composition in the mouse blastocyst. Nat Commun 7:13463. https://doi.org/10.1038/ncomms13463

22. Fischer SC, Corujo-Simon E, Lilao-Garzon J, Stelzer EHK, Muñoz-Descalzo S (2020) The transition from local to global patterns governs the differentiation of mouse blastocysts. PLoS One 15(5). https://doi.org/10.1371/journal.pone.0233030

23. Nichols J, Silva J, Roode M, Smith A (2009) Suppression of Erk signalling promotes ground state pluripotency in the mouse embryo. Development 136(19):3215–3222. https://doi.org/10.1242/dev.038893

24. Saiz N, Mora-Bitrià L, Rahman S, et al (2020) Growth-factor-mediated coupling between lineage size and cell fate choice underlies robustness of mammalian development. Elife 9:1–38. https://doi.org/10.7554/ELIFE.56079

25. Yu Q, Li J, Dai C ling, et al (2020) Anesthesia with sevoflurane or isoflurane induces severe hypoglycemia in neonatal mice. PLoS One 15(4):e0231090. https://doi.org/10.1371/JOURNAL.PONE.0231090

26. Chan YK, Davis PF, Poppitt SD, et al (2012) Influence of tail versus cardiac sampling on blood glucose and lipid profiles in mice. Lab Anim 46:142–147. https://doi.org/10.1258/la.2011.011136

27. Horber FF, Krayer S, Miles J, Cryer P, Rehder K, Haymond MW (1990) Isoflurane and whole body leucine, glucose, and fatty acid metabolism in dogs. Anesthesiology 73(1):82–92. https://doi.org/10.1097/00000542-199007000-00013

28. Thoolen B, Maronpot RR, Harada T, et al (2010) Proliferative and nonproliferative lesions of the rat and mouse hepatobiliary system. Toxicol Pathol 38(7 SUPPL.). https://doi.org/10.1177/0192623310386499

29. Tessitore A, Cicciarelli G, Del Vecchio F, et al (2016) MicroRNA expression analysis in high fat diet-induced NAFLD-NASH-HCC progression: study on C57BL/6J mice. BMC Cancer 16(1). https://doi.org/10.1186/S12885-015-2007-1

30. Hall WC, Ganaway JR, Rao GN, et al (1992) Histopathologic observations in weanling B6C3F1 mice and F344/N rats and their adult parental strains. Toxicol Pathol 20(2):146–154. https://doi.org/10.1177/019262339202000202

31. Barthold SW, Griffey SM, Percy DH. Pathology of Laboratory Rodents and Rabbits 4th Ed. Wiley-Blackwell, Ames. 2016.

32. Willard-Mack CL, Elmore SA, Hall WC, et al (2019) Nonproliferative and Proliferative Lesions of the Rat and Mouse Hematolymphoid System. Toxicol Pathol 47(6):665. https://doi.org/10.1177/0192623319867053

33. Picklo MJ, Idso J, Seeger DR, Aukema HM, Murphy EJ (2017) Comparative effects of high oleic acid vs high mixed saturated fatty acid obesogenic diets upon PUFA metabolism in mice. Prostaglandins, Leukot Essent Fat Acids 119:25–37. https://doi.org/10.1016/J.PLEFA.2017.03.001

34. Soleimanzad H, Montaner M, Ternier G, et al (2021) Obesity in Midlife Hampers Resting and Sensory-Evoked Cerebral Blood Flow in Mice. Obesity 29(1):150–158. https://doi.org/10.1002/OBY.23051

35. Bentham J, Di Cesare M, Bilano V, et al (2017) Worldwide trends in body-mass index, underweight, overweight, and obesity from 1975 to 2016: a pooled analysis of 2416 population-based measurement studies in 128·9 million children, adolescents, and adults. Lancet 390(10113):2627–2642. https://doi.org/10.1016/S0140-6736(17)32129-3

36. Life span as a biomarker. https://www.jax.org/research-and-faculty/research-labs/the-harrison-lab/gerontology/life-span-as-a-biomarker. Accessed 31 Mar 2022

37. Dutta S, Sengupta P (2016) Men and mice: Relating their ages. Life Sci 152:244–248. https://doi.org/10.1016/J.LFS.2015.10.025

38. Chaix A, Deota S, Bhardwaj R, Lin T, Panda S (2021) Sex- and age-dependent outcomes of 9-hour time-restricted feeding of a Western high-fat high-sucrose diet in C57BL/6J mice. Cell Rep 36(7):109543. https://doi.org/10.1016/J.CELREP.2021.109543

39. Brinton RD (2012) Minireview: Translational animal models of human menopause: Challenges and emerging opportunities. Endocrinology 153(8):3571–3578. https://doi.org/10.1210/en.2012-1340

40. Ozekinci M, Seven A, Olgan S, et al (2015) Does obesity have detrimental effects on IVF treatment outcomes? BMC Womens Health 15(1). https://doi.org/10.1186/S12905-015-0223-0

41. Bellver J, Ayllón Y, Ferrando M, et al (2010) Female obesity impairs in vitro fertilization outcome without affecting embryo quality. Fertil Steril 93(2):447–454. https://doi.org/10.1016/J.FERTNSTERT.2008.12.032

42. Qi L, Liu YP, Wang SM, et al (2022) Abnormal BMI in Male and/or Female Partners Are Deleterious for Embryonic Development and Pregnancy Outcome During ART Process: A Retrospective Study. Front Endocrinol (Lausanne) 13:517. https://doi.org/10.3389/FENDO.2022.856667/BIBTEX

43. Larsen MD, Jensen DM, Fedder J, Jølving LR, Nørgård BM (2020) Live-born children after assisted reproduction in women with type 1 diabetes and type 2 diabetes: a nationwide cohort study. Diabetologia 1–9. https://doi.org/10.1007/s00125-020-05193-6

44. Mattsson K, Nilsson-Condori E, Elmerstig E, et al (2021) Fertility outcomes in women with pre-existing type 2 diabetes—a prospective cohort study. Fertil Steril 116(2):505–513. https://doi.org/10.1016/j.fertnstert.2021.02.009

45. Miao X, Cui W (2022) Berberine alleviates LPS-induced apoptosis, oxidation, and skewed lineages during mouse preimplantation development. Biol Reprod. https://doi.org/10.1093/BIOLRE/IOAC002

46. Chiu YH, Chen H (2016) GATA3 inhibits GCM1 activity and trophoblast cell invasion. Sci Rep 6. https://doi.org/10.1038/SREP21630

47. Davalli P, Mitic T, Caporali A, Lauriola A, D’Arca D (2016) ROS, Cell Senescence, and Novel Molecular Mechanisms in Aging and Age-Related Diseases. Oxid Med Cell Longev 2016. https://doi.org/10.1155/2016/3565127

48. Fernández-Sánchez A, Madrigal-Santillán E, Bautista M, et al (2011) Inflammation, oxidative stress, and obesity. Int J Mol Sci 12(5):3117–3132. https://doi.org/10.3390/IJMS12053117

49. Jawerbaum A, White V (2010) Animal models in diabetes and pregnancy. Endocr Rev 31(5):680–701. https://doi.org/10.1210/er.2009-0038

50. Rehman K, Akash MSH (2017) Mechanism of Generation of Oxidative Stress and Pathophysiology of Type 2 Diabetes Mellitus: How Are They Interlinked? J Cell Biochem 118(11):3577–3585. https://doi.org/10.1002/JCB.26097

